# Immunological imprinting shapes the cross-reactive antibody responses to the KP.2 and LP.8.1 vaccine doses

**DOI:** 10.64898/2026.05.13.725047

**Authors:** Sanjeev Kumar, Lilin Lai, Madison L. Ellis, Anamika Patel, Devyani Joshi, Jacob Vander Velden, Jana Ziad Abu Faraj, Sonia T. Wimalasena, Ramitha R. Pallavi, Jad Iriss, Kareem Bechnak, Sri Edupuganti, Nadine Rouphael, Eric A. Ortlund, Alberto Moreno, Vineet D. Menachery, Veronika I. Zarnitsyna, Jens Wrammert, Mehul S. Suthar

## Abstract

The emergence of the SARS-CoV-2 Omicron BA.2.86 subvariant, a lineage derived from the BA.2 strain, led to the 2024-2025 COVID-19 vaccine update to include KP.2 or related JN.1-lineage spike antigens. We evaluated the magnitude, breadth, and durability of humoral immune responses following a single KP.2 vaccine dose in a longitudinal cohort of 21 individuals up to six months. KP.2 vaccination increased spike-specific binding and neutralizing antibodies against the ancestral WA1 strain, alongside the BA.5, XBB.1.5, and KP.2 variants. Power law modeling estimated half-lives for WA1- and KP.2-specific IgG responses at 770 and 248 days, respectively. Additionally, the KP.2 dose increased IgG1 and IgG4 subclasses more than IgG2 and IgG3 responses to both spike proteins. Serum depletion experiments using WA1 or KP.2 proteins demonstrated most vaccine-elicited antibodies were cross-reactive. Consequently, KP.2 vaccine-induced antibodies retained broad neutralizing activity against recently circulating Omicron subvariants (BA.2.86, KP.3.1.1, XEC, LP.8.1, LF.7, XFG.3.12, PQ.1, BA.3.2.1, and RE.2). Using a live virus neutralization assay, XFG.3.12 showed the greatest reduction in neutralizing titers relative to KP.2 (4.2-fold). In a small subset, an LP.8.1 vaccine dose increased neutralizing activity against the matched variant while maintaining WA1 and KP.2 cross-reactivity, but only modestly increased antibodies to divergent variants BA.3.2.1 and RE.2. Ultimately, these data indicate the KP.2 mRNA vaccine generates durable, cross-reactive responses against current Omicron subvariants. However, ongoing spike evolution impacts neutralization of emerging lineages, highlighting the need for continued viral monitoring and timely vaccine updates.

**IMPORTANCE:** SARS-CoV-2 continues to evolve, raising ongoing concerns about how well updated vaccines protect against emerging variants. This study evaluates antibody responses after KP.2 spike mRNA vaccine dose and shows that a single dose induces durable and broadly cross-reactive immunity against both earlier strains and recently circulating Omicron subvariants. Despite this breadth, reduced neutralizing activity against certain emerging variants indicates that ongoing antigenic changes can impact vaccine induced antibody effectiveness. These findings provide insight into how current vaccines perform over time and highlight the need to track viral evolution and update vaccine antigens to maintain broad protection against severe disease, hospitalization, and death.

## INTRODUCTION

Vaccination remains a key strategy for limiting severe disease caused by severe acute respiratory syndrome coronavirus 2 (SARS-CoV-2) (1-3). Current vaccines are largely designed around the viral spike glycoprotein, which mediates attachment to host cells through interaction with the host angiotensin-converting enzyme 2 (ACE2) which serves as the primary target of neutralizing antibodies (4). Continual viral transmission and global cycles of viral emergence and spread have resulted in the accumulation of mutations within the spike protein, which continue to threaten the effectiveness of SARS-CoV-2 vaccines and mononclonal antibody therapeutics (5-8). To address this, the composition of the vaccine dose is periodically updated to reflect the antigenic changes in circulating variants (9-13). Following the initial ancestral WA1 strain based three vaccine doses, bivalent formulations incorporating both ancestral WA1 and Omicron spike proteins BA.4/BA.5 were introduced to broaden immune responses, followed by monovalent vaccines based on the Omicron sublineage XBB.1.5 (14-17). Continued viral evolution has led to further updates, including vaccines based on the JN1-lineage spike antigen for the 2024-2025 immunization season (9).

The antigenic landscape of SARS-CoV-2 has become increasingly complex, particularly with the emergence of Omicron variants that contain extensive mutations in the spike protein (18). These mutations alter key neutralizing epitopes, enabling escape from antibodies elicited by prior infection or vaccination (19, 20). Antigenic mapping studies have shown that recent Omicron subvariants occupy distinct positions relative to earlier strains, reflecting substantial divergence in antibody recognition (7, 21, 22). The BA.2.86 lineage represents a notable evolutionary step, characterized by a large number of spike mutations compared to earlier BA.2 derived viral strains (20). Descendants of this lineage, including KP.2, have spread globally and have contributed to waves of infection (21, 23). The antigenic distance between these variants and prior vaccine strains raised concerns regarding the effectiveness of existing immunity and prompted the selection of KP.2 spike protein for updated vaccines (7). Understanding how these vaccine perform against both matched and mismatched variants is essential for guiding future vaccine design and vaccination strategies.

An important determinant of vaccine performance is the persistence of protective antibody responses over time. Circulating antibody levels typically decline following vaccination, although the rate of decay and the duration of detectable neutralizing activity can vary depending on prior exposure history, vaccine platform, and antigen matching (1, 11, 24-26). In addition, the breadth of antibody responses across antigenically distinct variants is critical, particularly in the context of rapidly evolving viral strains (8, 22). It has been observed that repeated antigen exposure through infection or vaccination can shape the antibody repertoire, often leading to cross-reactive responses that recognize conserved regions of the spike protein (27-29). While such responses may confer broad protection, they can also reflect immune imprinting, where prior antigenic exposures influence the specificity of subsequent responses (29-31). This balance between breadth and specificity is especially relevant for updated vaccines targeting divergent variants.

In this study, we assessed the durability and breadth of antibody responses following immunization with a KP.2 spike based SARS-CoV-2 mRNA vaccine. Using a longitudinal cohort, we measured spike-specific binding antibodies, IgG subclass responses, and live virus neutralization against both ancestral and recently circulating variants over a period of six-months. We further evaluated cross-reactivity through serum depletion experiments to determine the contribution of shared versus variant-specific antibody populations. Our findings show that KP.2 spike mRNA vaccine dose induces durable and broadly cross-reactive antibody responses, with sustained neutralizing activity against multiple Omicron subvariants. Further, reduced neutralization of certain contemporary SARS-CoV-2 Omicron subvariants highlights ongoing antigenic drift and the challenges it poses for vaccine design. These results provide insight into the performance of updated SARS-CoV-2 vaccines and inform strategies for future immunogen or vaccine antigen selection.

## RESULTS

### A KP.2 spike vaccine dose preferentially improves spike binding and neutralizing antibodies against the KP.2 variant

A prospective longitudinal study was conducted to evaluate immune responses induced by a monovalent KP.2 vaccine dose in a cohort of 21 participants. Blood samples were collected before and subsequently at 1-, 3-, and 6-months following vaccination from participants enrolled at Emory University (Emory Vaccine Center and Emory Hope Clinic). Our study aimed to assess the magnitude, breadth, and persistence of vaccine-induced binding and neutralizing antibodies, over a six-month period after KP.2 spike vaccine dose (**Table S1**). First, to understand the KP.2 variant diversity relative to other key SARS-CoV-2 variants, we performed phylogenetic and spike structural analyses (**Fig. 1**). We found that KP.2 variant clusters closely with the BA.2.86 variant and is antigenically distinct from ancestral WA1 and earlier Omicron variants BA.5 and XBB.1.5 (**Fig. 1A**). Structural mapping of spike mutations showed both shared and unique substitutions across BA.5, XBB.1.5, and KP.2, with several changes localized to the receptor binding domain (**Fig. 1B-E**).

**Fig. 1.**
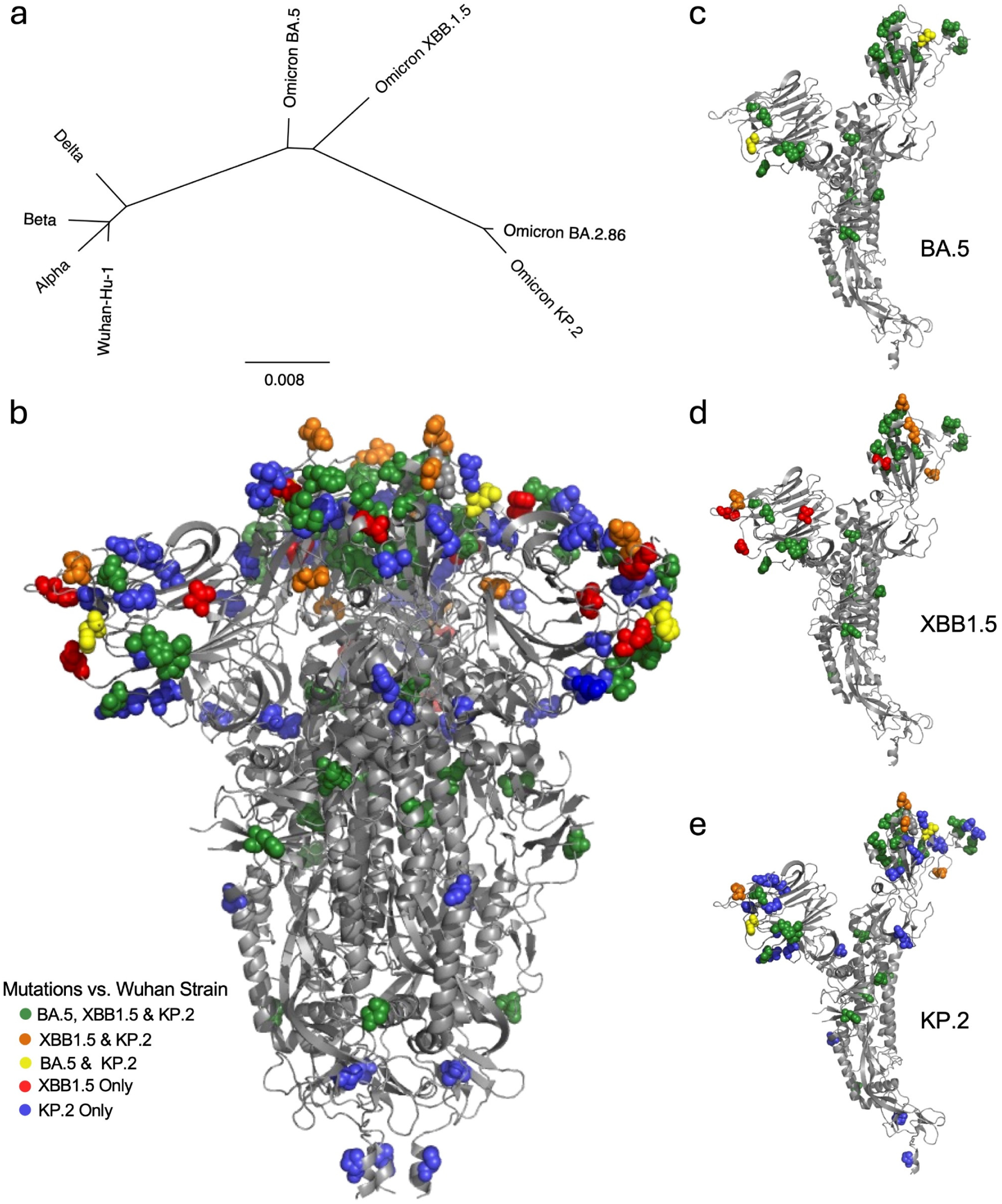
Phylogenetic and structural presentation of SARS-CoV-2 spike variants used in mRNA vaccines. **(A)** Phylogenetic tree of SARS-CoV-2 spike sequences showing the evolutionary relationship between the ancestral Wuhan-Hu-1 strain and SARS-CoV-2 variants, including Alpha, Beta, Delta, and Omicron sublineages BA.5, XBB.1.5, BA.2.86 and KP.2. **(B)** Structural representation of the trimeric spike protein highlighting shared and unique mutations in BA.5, XBB.1.5, and KP.2. Mutations are color-coded to indicate overlap between variants and variant-specific substitutions. **(C-E)** Monomeric spike structures of **(C)** BA.5, **(D)** XBB.1.5, and **(E)** KP.2, with mutations mapped onto the spike protein.

Next, the plasma samples of all the study participants were assessed to quantify the neutralizing antibody titers against WA1, BA.5, XBB.1.5 and KP.2 using a live SARS-CoV-2 focus reduction neutralization test (FRNT), and FRNT_50_ geometric mean titers (GMTs) were calculated. FRNT_50_ titers undetectable below 1:20 dilution were assigned a value of 20 and marked as non-responders. We found that all study samples collected prior to KP.2 vaccination exhibit detectable FRNT_50_ neutralizing antibodies titer against WA1 and BA.5 viruses, whereas, reduced number (86%) of individuals had neutralizing antibodies against both XBB.1.5 and KP.2 Omicron subvariants (**Fig. 2A**). The neutralizing antibody titers were significantly lower against BA.5 (*p* = 0.0135), XBB.1.5 (*p* < 0.0001), and KP.2 (*p* < 0.0001) compared to WA1 with a difference of FRNT_50_ GMT titers 4.1- for BA.5, >10.3- for XBB.1.5, and >10.7-fold for KP.2. The FRNT_50_ GMTs were 715 for WA1, 174 for BA.5, 69.6 for XBB.1.5, and 67 for KP.2 (**Fig. 2A, left panel**). At 1-month post-vaccination, FRNT_50_ GMTs were increased against all four viruses, with titer of 1053 for WA1, 400 for BA.5, 181 for XBB.1.5 and 456 for KP.2 variant with a difference of 2.6- for BA.5, >5.8- for XBB.1.5, and >2.3-fold for KP.2, relative to WA1. At this time point 95% of the participants had detectable neutralizing antibodies titers for both XBB.1.5 and KP.2 variants (**Fig. 2A, second panel from left**). Between 1 and 3 months, neutralizing titers declined modestly with a difference of 3.1- for BA.5, > 6.8- for XBB.1.5, and >3-fold for KP.2, relative to WA1 (**Fig. 2A, third panel from left**), followed by a slower rate of decline between 3 and 6 months with a difference of 3.2- for BA.5, >6.8- for XBB.1.5, and >3.1-fold for KP.2, relative to WA1 (**Fig. 2A, right panel**).

**Fig. 2.**
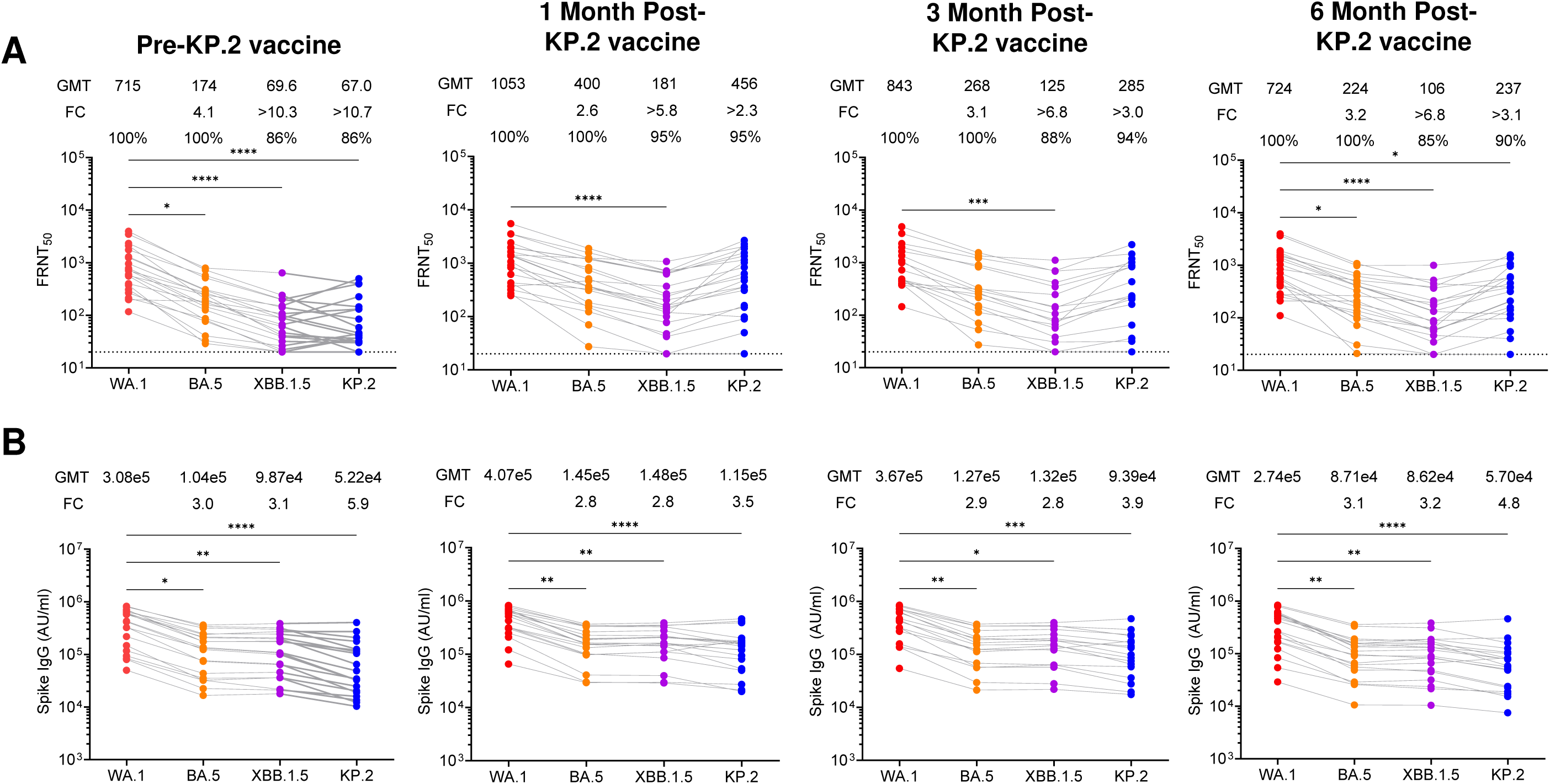
Neutralizing and binding antibody responses following KP.2 spike mRNA vaccination against WA1 and Omicron subvariants. **(A)** Neutralizing antibody responses measured by using live-virus based focus reduction neutralization test (FRNT_50_) assays against WA1, BA.5, XBB.1.5, and KP.2 at pre-vaccination (Pre), 1 month (1M), 3 months (3M), and 6 months (6M) after KP.2 vaccination (n = 21). FRNT_50_ geometric mean titers (GMTs) are indicated above each group along with the ratio (FC) of the neutralization GMT against WA1 and the Omicron subvariants. **(B)** Spike-specific IgG binding titers measured by electrochemiluminescence assay against WA1, BA.5, XBB.1.5, and KP.2 at the same time points. Titers are reported as arbitrary units per milliliter (AU/mL). Binding GMTs are indicated above each group along with the ratio (FC) of the binding GMT against WA1 and the Omicron subvariants. For both panels, each point represents an individual, and lines connect matched samples across time points. Statistical comparisons were performed using Kruskal-Wallis test with Dunn’s multiple comparisons test. *p < 0.05, **p < 0.01, ***p < 0.001, ****p < 0.0001.

We next measured Spike-specific IgG binding titers using an electrochemiluminescence assay (**Fig. 2B**). We found that the binding antibody titers were significantly lower against BA.5 (*p* = 0.0181), XBB.1.5 (*p* = 0.0098), and KP.2 (*p* < 0.0001) compared to WA1 with a difference of 3.0- for BA.5, 3.1- for XBB.1.5, and 5.9-fold for KP.2. The GMTs were 3.08x10^5^ for WA1, 1.04x10^5^ for BA.5, 9.87x10^4^ for XBB.1.5, and 5.22x10^4^ for KP.2 (**Fig. 2B, left panel**). At 1-month post-vaccination, the binding antibody titers were significantly lower against BA.5 (*p* = 0.0010), XBB.1.5 (*p* = 0.0016), and KP.2 (*p* < 0.0001) relative to WA1 with a difference of 2.8- for BA.5, 2.8- for XBB.1.5, and 3.5-fold for KP.2. The GMTs were 4.07x10^5^ for WA1, 1.45x10^5^ for BA.5, 1.48x10^5^ for XBB.1.5, and 1.15x10^5^ for KP.2 (**Fig. 2B, second panel from left**). At 3 months, neutralizing titers declined modestly with a difference of 2.9-fold (*p* = 0.0099) for BA.5, 2.8-fold (*p* = 0.0166) for XBB.1.5, 3.9-fold (*p* = 0.0005) for KP.2 compared to WA1 (**Fig. 2B, third panel from left**), followed by a slower rate of decline at 6 months with a difference 3.1-fold (*p* = 0.0054) for BA.5, 3.2-fold (*p* = 0.0043) for XBB.1.5, and 4.8-fold (*p* < 0.0001) for KP.2 relative to WA1 (**Fig. 2B, right panel**). Overall, these results show that a single KP.2 spike mRNA vaccine dose induced robust neutralzing and binding antibody response to all the viruses tested including the ancestral WA1 strain and KP.2 Omicron subvariant with stable neutralizing and binding titers observed up to six months.

### KP.2 spike mRNA vaccine induces durable antibody responses

We performed longitudinal analysis of neutralizing antibody titers in a subset of randomly selected individuals (n = 10) and followed them for up to six months after the KP.2 vaccine dose (**Fig. 2**). These individuals showed negligible change in antibody titers to the SARS-CoV-2 nucleocapsid protein (**Fig. S1**) suggesting that these individuals did not get any SARS-CoV-2 breakthrough infection during the 6 months following KP.2 vaccination. The neutralizing antibody responses against WA.1, BA.5, XBB.1.5, and KP.2 variants increased substantially one month after vaccination and subsequently declined gradually over time. For the ancestral WA.1 strain, the geometric mean neutralization titer (GMT) increased from 716 at baseline to 1089 at 1 month, followed by a gradual decline to 789 at 3 months and 632 at 6 months (**Fig. 3A**). A similar pattern was observed for BA.5, where GMT increased from 181 at baseline to 463 at 1 month, then decreased to 253 at 3 months and 208 at 6 months (**Fig. 3B**). Neutralization of XBB.1.5 also increased after vaccination, with GMT increasing from 70.1 pre-vaccination to 189 at 1 month, followed by a decline to 106 at 3 months and 94 at 6 months (**Fig. 3C**). Similarly, neutralizing antibody titers against the vaccine-matched KP.2 variant increased markedly from 73.6 pre-vaccination to 494 at 1 month, followed by gradual waning to 222 at 3 months and 173 at 6 months (**Fig. 3D**). Despite this decline over time, neutralizing titers remained above pre-vaccination baseline levels at 6 months for all variants tested, indicating sustained neutralizing activity.

**Fig. 3.**
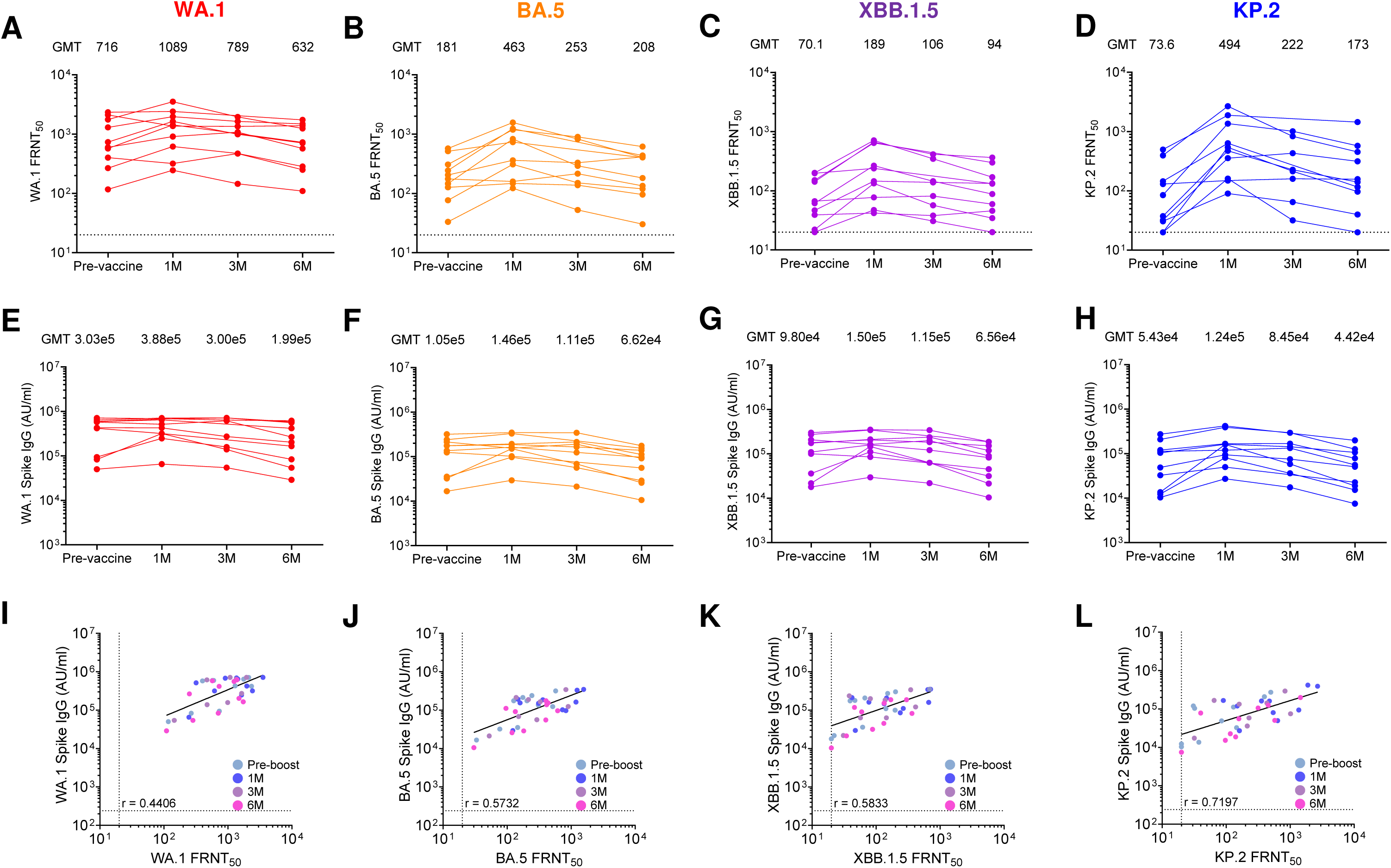
Durability and correlation of neutralizing and binding antibody responses following KP.2 spike mRNA vaccination. (A-D) These are the longitudinal neutralizing antibody titers (FRNT_50_) against **(A)** WA1, **(B)** BA.5, **(C)** XBB.1.5, and **(D)** KP.2 measured at pre-vaccination (Pre), 1 month (1M), 3 months (3M), and 6 months (6M) post-vaccination. FRNT_50_ geometric mean titers (GMTs) are indicated above each group. Each point represents an individual, with lines connecting longitudinal samples. **(E-H)** These are the corresponding spike-binding IgG titers as AU/mL for **(E)** WA1, **(F)** BA.5, **(G)** XBB.1.5, and **(H)** KP.2 measured over time. Binding GMTs are indicated above each group. Each point represents an individual, with lines connecting longitudinal samples. **(I-L)** Correlation between spike-binding IgG titers and neutralizing antibody titers (FRNT_50_) for **(I)** WA1, **(J)** BA.5, **(K)** XBB.1.5, and **(L)** KP.2 across all time points are shown in which each samples dot is shown in different colors for each time point. Spearman correlation coefficients (r) are shown in panels **I-L**.

We next examined spike-binding IgG antibody responses over the same time period of six-months. Spike binding antibody titers showed greater stability than neutralizing titers across all variants. For WA.1 spike, GMT values were 3.03x10^5^ AU/ml at baseline, increasing modestly to 3.88x10^5^ AU/ml at 1 month, and remaining relatively stable at 3.00x10^5^ AU/ml at 3 months and 1.99x10^5^ AU/ml at 6 months (**Fig. 3E**). BA.5 spike-binding IgG showed a similar profile with GMT values of 1.05x10^5^ AU/ml pre-vaccination, increasing to 1.46x10^5^ AU/ml at 1 month, and remaining relatively stable at 1.11x10^5^ AU/ml at 3 months and 6.62x10^4^ AU/ml at 6 months (**Fig. 3F**). For XBB.1.5 spike, GMT values were 9.80x10^4^ AU/ml at baseline, 1.50x10⁵ AU/ml at 1 month, 1.15x10^5^ AU/ml at 3 months, and 6.56x10^4^ AU/ml at 6 months (**Fig. 3G**). Similarly, KP.2 spike-binding IgG increased from 5.43x10^4^ AU/ml pre-vaccination to 1.24x10^5^ AU/ml at 1 month, followed by a modest decline to 8.45x10^4^ AU/ml at 3 months and 4.42x10^4^ AU/ml at 6 months (**Fig. 3H**). Overall, spike-binding antibody titers remained relatively stable across the six-month period, with only modest declines after the peak response was observed.

To determine the relationship between binding and neutralizing antibody responses, we next performed correlation analysis between spike-binding IgG levels and FRNT_50_ neutralization titers. We observed positive correlations for all variants tested (**Fig. 3I-L**), with Spearman correlation coefficients of r = 0.44 for WA.1, r = 0.57 for BA.5, r = 0.58 for XBB.1.5, and r = 0.72 for KP.2, indicating that higher spike-binding antibody levels were associated with stronger neutralizing activity.

To estimate antibody durability, we modeled decay rates using the power law and exponential decay models (**Fig. 4**). For WA.1 spike-binding IgG antibodies, the power law decay model estimated a half-life (t_1/2_) of 206 days, whereas the exponential decay model estimated a longer half-life of 770 days (**Fig. 4A**). For BA.5 spike-binding antibodies, the estimated half-lives were 120 days by the power law model and 248 days by the exponential model (**Fig. 4B**). In the case of XBB.1.5 spike-binding antibodies, the estimated half-life was 217 days by the power law model and 1359 days by the exponential model (**Fig. 4C**). For KP.2 spike-binding antibodies, the decay half-lives were 114 days by the power law model and 390 days by the exponential model (**Fig. 4D**). Overall, these modeling data indicate that spike-binding antibodies elicited by the KP.2 vaccine persist for several months with gradual decay kinetics. Together, these results demonstrate that KP.2 mRNA vaccination induces robust neutralizing and spike-binding antibody responses that peak one month after vaccination and remain detectable for at least six months. The relatively slow decay kinetics and sustained binding antibody levels suggest that KP.2 vaccination generates durable humoral immunity against both ancestral and contemporary SARS-CoV-2 Omicron subvariants.

**Fig. 4.**
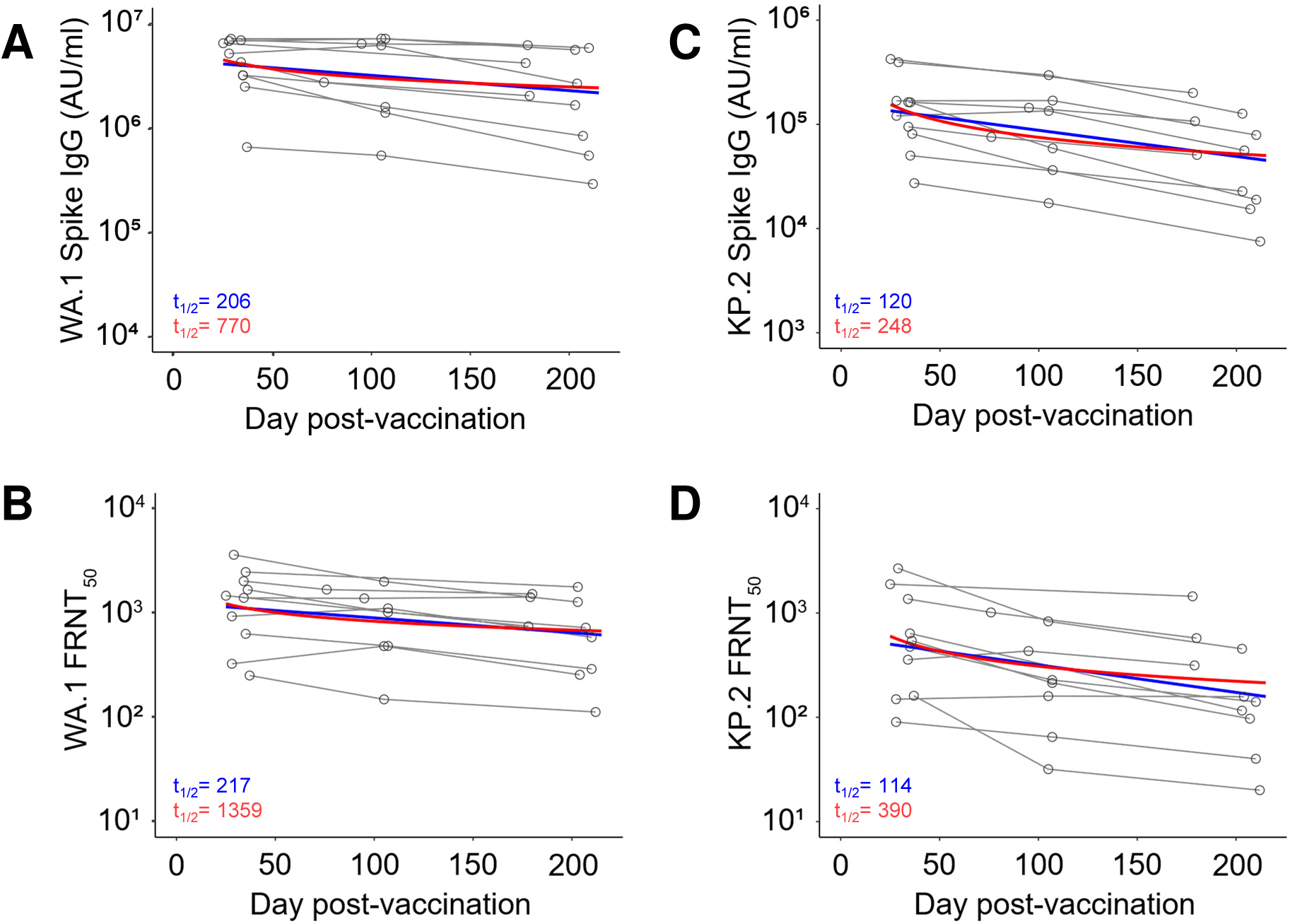
Estimation of antibody decay following KP.2 vaccination. Longitudinal data for the ten individuals is represented by grey open circles connected by lines for each individual. Shown are the **(A)** WA1 binding data; **(B)** WA1 neutralization data; **(C)** KP.2 binding data; and **(D)** KP.2 neutralization data. Also shown are two decay curves and their corresponding estimated half-lives, derived from fitting two different models: the exponential (blue), and power-law (red) models. For the power-law decay model, the half-life was estimated at day 90 post-vaccination. The connecting lines between the variants in panels **(A-D)** represent matched serum samples.

### KP.2 mRNA vaccination elicits durable IgG subclass spike protein-binding antibodies

Our previous study found that the XBB.1.5 vaccine dose induced IgG1, Ig3, and IgG4 subclass antibodies that were differentially sustained through 6 months post-vaccine dose (11). We next assessed the durability of IgG subclass antibody binding to the ancestral WA1 and KP.2 spike proteins following KP.2 vaccination using electrochemiluminescence assays. Normalized geometric mean titers (nGMTs) calculated by subtracting background binding measured in pre-pandemic control samples, were used to quantify subclass-specific responses across the longitudinal time points. For IgG1, before vaccination we observed a 6.1-fold lower binding to the KP.2 spike compared with WA1 (WA1 nGMT 121, KP.2 nGMT 19.9). One month after KP.2 vaccination, IgG1 binding increased substantially to both spike proteins, with WA1 reaching an nGMT of 218 and KP.2 reaching 73.7, corresponding to a 2.9-fold difference between the two antigens (**Fig. 5A**). In the longitudinal cohort, the binding titers were increased from baseline to 1 month with nGMT of 137 to 203 for WA1 and 24.6 to 77.6 for KP.2 (**Fig. 5B and 5C**), indicating a stronger boosting of antibodies directed toward the vaccine-matched antigen. IgG1 titers gradually declined after the peak response, with a larger reduction occurring between 1 and 3 months than between 3 and 6 months post-vaccination. Despite this decline, all samples maintained IgG1 binding above background levels throughout the follow-up period.

**Fig. 5.**
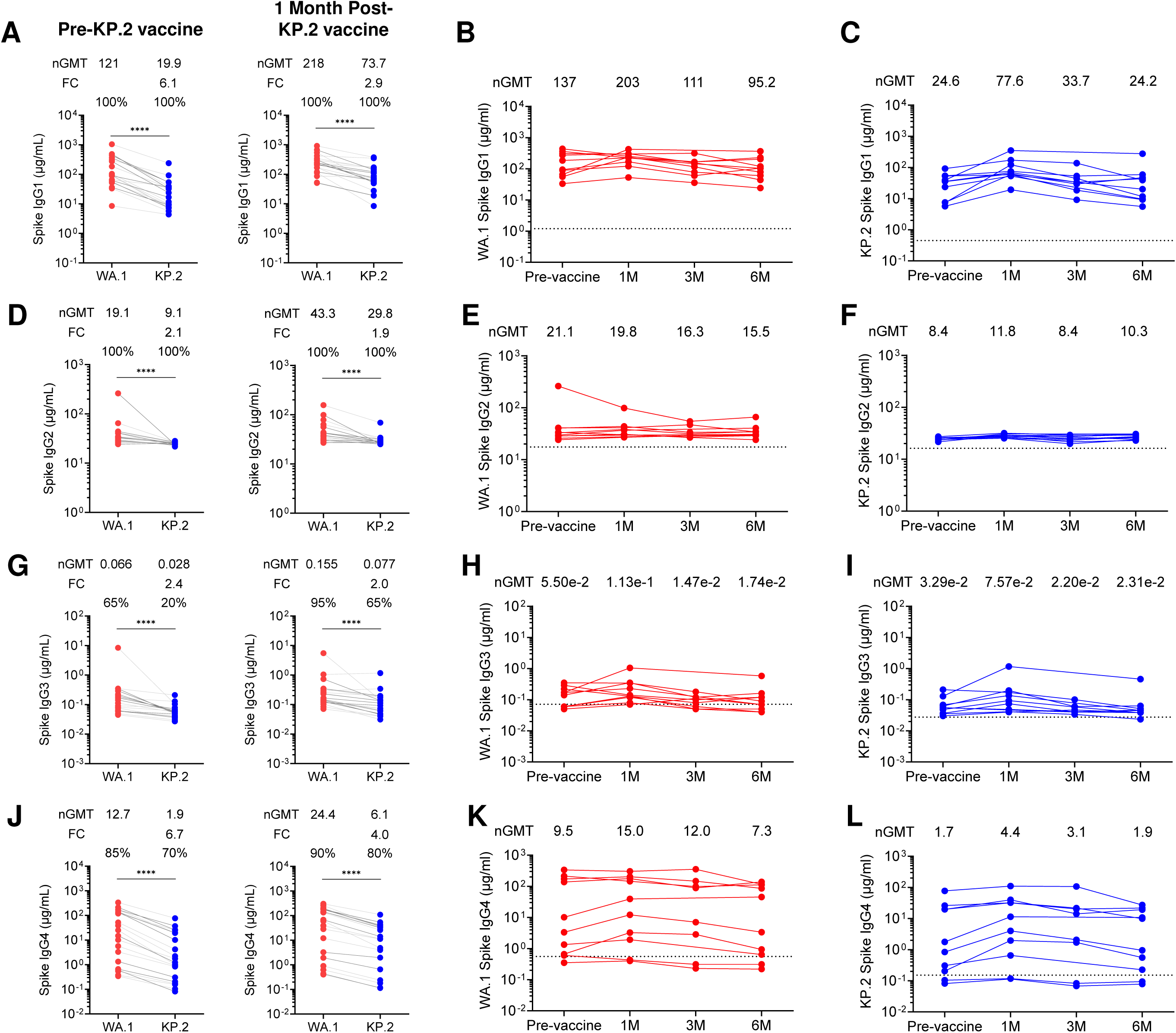
IgG subclass responses to SARS-CoV-2 spike following KP.2 mRNA vaccination. The IgG antibody subclass binding titers to the SARS-CoV-2 spike proteins of WA1 and KP.2 strains were determined by an electrochemiluminescence based assay by mesoscale discovery assay (MSD). Each dot represents the titer obtained for each individual in micrograms per ml (µg/mL) which was extrapolated from standard curves generated from titer of monoclonal antibodies specific for each IgG subclass. Left panels (**A**) IgG1, **(D)** IgG2, **(G)** IgG3 and **(J)** IgG4 show subclass-specific binding to WA1 and KP.2 spike proteins at pre-vaccination and 1-month post-vaccination for all 21 participants. Binding GMTs are indicated above each group along with the ratio (FC) of the binding GMT against WA1 and KP.2 omicron subvariant. Statistical comparisons were performed using Wilcoxon matched-pairs signed rank test for paired comparisons. ****p < 0.0001. Middle panels show longitudinal binding responses specific for (**B**) IgG1, **(E)** IgG2, **(H)** IgG3 and **(K)** IgG4 subclass to WA1 spike protein in a subset of individuals (n=10) through 6 months. Binding GMTs are indicated above each group. Right panels show longitudinal binding responses specific for (**C**) IgG1, **(F)** IgG2, **(I)** IgG3 and **(L)** IgG4 subclass to KP.2 spike protein in a subset of individuals (n=10) through 6 months. Binding GMTs are indicated above each group.

For IgG2, prior to vaccination we observed a 2.1-fold lower binding to KP.2 compared with WA1 (WA1 nGMT 19.1; KP.2 nGMT 9.1). At one month following vaccination, IgG2 binding increased to WA1 nGMT 43.3 and KP.2 nGMT 29.8, corresponding to a 1.9-fold difference between the two spike proteins (**Fig. 5D**). In the longitudinal cohort, the binding titers were decreased from baseline to 1 month with nGMT of 21.1 to 19.8 for WA1 and increased from 8.4 to 11.8 for KP.2 (**Fig. 5E and 5F**). Considerable inter-individual variability was observed for this subclass. Over time, IgG2 titers steadily decreased between the 1-, 3-, and 6-month time points. By 6 months, several KP.2-specific IgG2 responses did not change much and remained detectable in most individuals.

For IgG3, baseline antibody binding to both spike proteins was low, with nGMT values of 0.066 for WA1 and 0.028 for KP.2, representing a 2.4-fold reduction for KP.2 relative to WA1. Following vaccination, IgG3 binding increased to 0.155 for WA1 and 0.077 for KP.2 at one month, corresponding to a 2-fold difference between the two spike proteins (**Fig. 5G**). In the longitudinal cohort, the binding titers were increased from baseline to 1 month with nGMT of 5.50E-02 to 1.13E-01 for WA1 and modestly increased from 3.29E-02 to 7.57E-02 for KP.2 (**Fig. 5H-I**). Similar to IgG1, IgG3 titers declined after the peak response, with the largest decrease occurring between 1 and 3 months, followed by a slower decline between 3 and 6 months.

For IgG4, baseline samples showed lower binding to KP.2 relative to WA1, with nGMT values of 12.7 for WA1 and 1.9 for KP.2, representing a 6.7-fold difference between the spike proteins. One month after vaccination, IgG4 binding increased to 24.4 for WA1 and 6.1 for KP.2, corresponding to a 4-fold reduction for KP.2 relative to WA1 (**Fig. 5J**). In the longitudinal cohort, the binding titers were increased from baseline to 1 month with nGMT of 9.5 to 15 for WA1 and 1.7 to 4.4 for KP.2 (**Fig. 5K-L**), indicating a greater vaccine-driven expansion of KP.2-directed IgG4 responses. IgG4 binding gradually declined between the 1-, 3-, and 6-month time points but remained detectable in most individuals throughout the observation period. Together, these findings demonstrate that KP.2 vaccination enhances spike-specific IgG subclass responses with distinct kinetics. The greatest increases were observed for IgG1, followed by IgG4, IgG3, and IgG2 and displayed greater variability across individuals. Overall, KP.2 vaccination induced subclass antibody responses that remained detectable for at least six months after immunization.

### Serum depletion with SARS-CoV-2 spike protein shows an immune-imprinted antibody response after KP.2 vaccination

To understand how prior exposures imprint antibody responses to the KP.2 vaccine, we performed spike protein depletion assays using magnetic beads conjugated with either ancestral WA1 or KP.2 spike proteins. Serum samples collected before vaccination and one month after vaccination from a subset of randomly selected participants (n = 10) were used for depletion experiments and also evaluated for residual spike-binding activity. When WA1 spike coupled beads were used to deplete serum antibodies, WA1 spike-binding reactivity was almost completely eliminated in both pre- and post-vaccination samples. Specifically, depletion resulted in a 146-fold significant reduction before vaccination and an 123-fold significant reduction after vaccination in WA1 spike-binding titers (**Fig. 6A, left panel**). In contrast, depletion with KP.2 spike protein produced a more modest reduction in WA1 spike-binding activity, with significant decreases of 2.4-fold before vaccination and 3.3-fold significant decrease after vaccination (**Fig. 6A, middle panel**). Quantification of antibody fractions showed that the proportion of WA1-reactive antibodies significantly declined from 47% before vaccination to 36% one month after vaccination (**Fig. 6A, right panel**), indicating a relative shift in antibody composition after immunization.

**Fig. 6.**
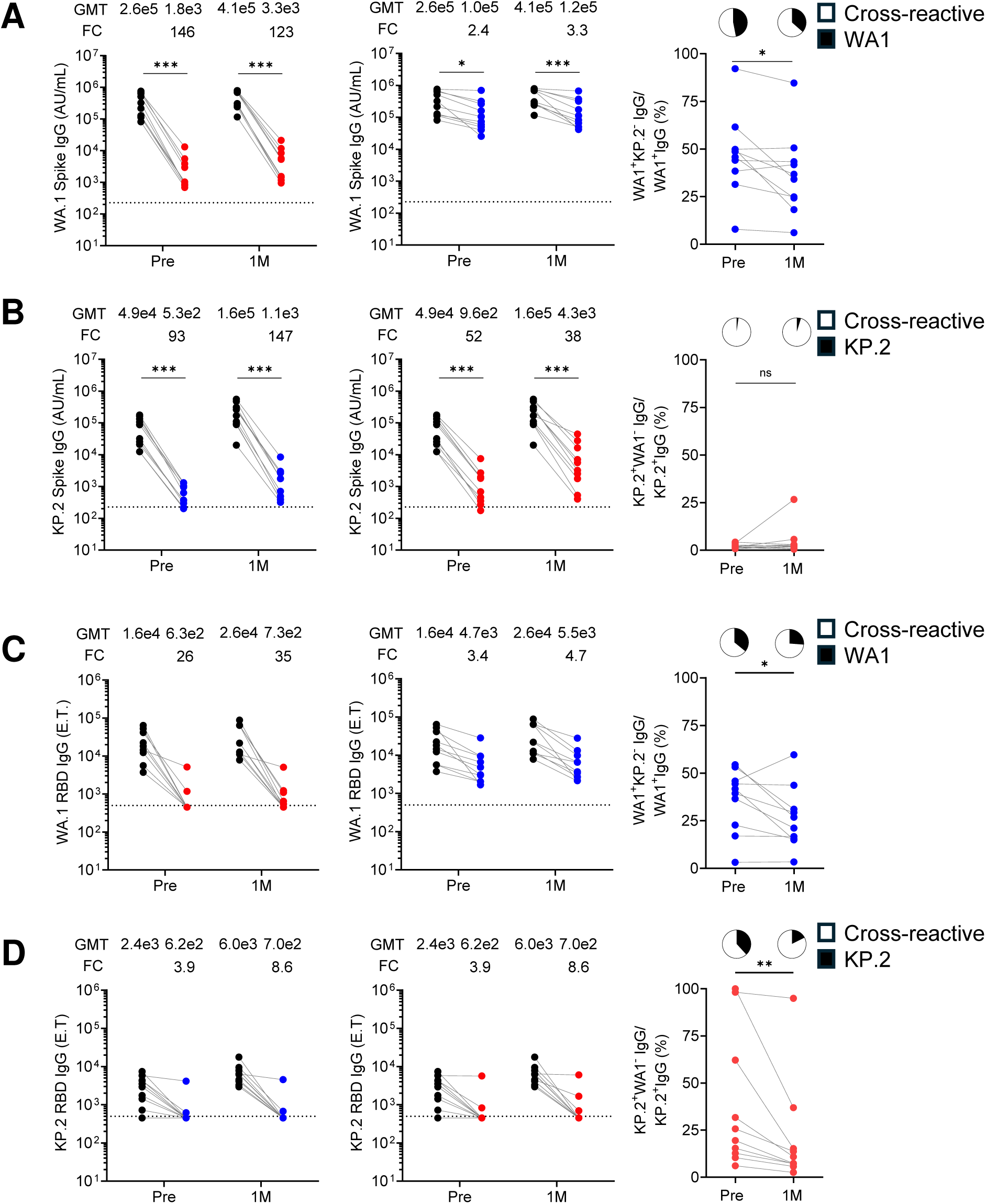
Serum depletion analysis reveals cross-reactive antibody responses following KP.2 vaccination. WA1 (red color) or KP.2 (blue color) spike trimeric protein depletion of serum samples from a subset of individuals (n=10) collected pre- and 1 month post-KP.2 vaccination was performed, followed by measurement of IgG binding to **(A)** WA1 spike proteins. **(B)** and KP.2 spike protein. Right panel in **(A)** represents the quantification of remaining WA1 cross-reactive antibodies, and right panel in **(B)** shows the quantification of remaining KP.2 cross-reactive antibodies fractions after depletion. Binding GMTs are indicated above each group along with the ratio (FC) of the binding GMT against WA1 and KP.2 omicron subvariant spike protein. **(C-D)** Shows WA1 (red color) or KP.2 (blue color) RBD protein depletion of serum samples from a subset of individuals (n=10) collected pre- and 1 month post-KP.2 vaccination was performed, followed by measurement of IgG binding to **(C)** WA1 RBD proteins. **(D)** and KP.2 RBD protein. Right panel in **(C)** represents the quantification of remaining cross-reactive and WA1 RBD-specific antibody fractions after depletion, and right panel in **(D)** shows the quantification of remaining cross-reactive and KP.2 RBD-specific antibody fractions after depletion. Binding GMTs are indicated above each group along with the ratio (FC) of the binding GMT against WA1 and KP.2 omicron subvariant RBD protein. Each point represents an individual sample. Statistical comparisons were performed using Wilcoxon matched-pairs test as appropriate. ns, not significant; *p < 0.05, **p < 0.01, ***p < 0.001.

Analogously, depletion using KP.2 spike conjugated beads resulted in a significant reduction in KP.2 spike-binding activity, with 93-fold and 147-fold decreases in pre- and post-vaccination samples, respectively (**Fig. 6B, left panel**). When serum samples were instead depleted using WA1 spike protein, KP.2 spike-binding titers were significantly reduced by 52-fold before vaccination and 38-fold after vaccination (**Fig. 6B, middle panel**). These findings indicate that the majority of antibodies recognizing the KP.2 spike protein are also cross-reactive with the ancestral spike antigen. A small fraction of KP.2-specific antibodies that were not removed by WA1 depletion increased modestly following vaccination, rising from 2.1% before vaccination to 4.8% at one month post-vaccination (**Fig. 6B, right panel**). This increase suggests the emergence of a limited pool of antibodies that preferentially recognize KP.2-specific epitopes after vaccination.

We further examined antibody binding directed to the receptor-binding domain (RBD) in the spike depleted samples. WA1 RBD-binding reactivity was reduced in both pre- and post-vaccination samples. Specifically, depletion resulted in a 26-fold reduction before vaccination and an 35-fold reduction after vaccination in WA1 RBD-binding titers (**Fig. 6C, left panel**). In contrast, depletion with KP.2 spike protein produced a more modest reduction in WA1 RBD-binding activity, with significant decreases of 3.4-fold before vaccination and 4.7-fold significant decrease after vaccination (**Fig. 6C, middle panel**). Quantification of antibody fractions showed that the proportion of WA1 RBD-specific antibodies significantly declined from 36% before vaccination to 26% one month after KP.2 vaccination (**Fig. 6C, right panel**), indicating a relative shift in antibody composition towards cross-reactive epitope after vaccination.

Depletion using KP.2 spike conjugated beads resulted in a modest reduction in KP.2 RBD-binding activity, with 3.9-fold and 8.6-fold decreases in pre- and post-vaccination samples, respectively (**Fig. 6D, left panel**). When serum samples were instead depleted using WA1 spike protein, KP.2 RBD-binding titers were significantly reduced by 3.9-fold before vaccination and 8.6-fold after vaccination (**Fig. 6D, middle panel**). These findings indicate that the majority of antibodies recognizing the KP.2 RBD protein are also cross-reactive with the ancestral spike antigen. A small fraction of KP.2 RBD-specific antibodies that were not removed by WA1 depletion decreased modestly following vaccination, declined from 39% before vaccination to 20% at one month post-vaccination (**Fig. 6B, right panel**). This decline suggests the emergence of a non-RBD pool of antibodies that preferentially recognize cross-reactive epitopes after vaccination.

Together, these results indicate that the antibody response induced by KP.2 vaccination is strongly influenced by previously established immune memory targeting conserved regions of the ancestral spike protein. Although vaccination modestly expands antibodies that recognize KP.2-spike specific epitopes, the overall response remains dominated by cross-reactive antibodies consistent with immune imprinting from prior SARS-CoV-2 exposures.

### KP.2 spike mRNA vaccination improves antibody neutralizing activity against contemporary Omicron variants

We evaluated the neutralizing activity against a panel of recently circulating Omicron subvariants, including BA.2.86, KP.3.1.1, XEC, LP.8.1.1, LF.7, XFG.3.12, and PQ.1, compared against KP.2 **(Fig. 7)**. At one month post-vaccination, neutralizing titers were detectable against all the variants. The FRNT_50_ GMTs were 354 for KP.2, 893 for BA.2.86, 163 for KP.3.1.1, 157 for XEC, 116 for LF.7, 85 for XFG.3.12, and 140 for PQ.1 **(Fig. 7A)**. With the exception of BA.2.86, which is a pre-cursor to the KP.2 variant, all of the Omicron subvariants showed reduced neutralizing antibody titers relative to KP.2. The neutralizing antibody titers for all of the Omicron subvairants continued to reduce through 6 months post-KP.2 vaccine dose, but remained detectable in the majority of study participants **(Fig. 7B)**. The reduction in neutralization relative to KP.2 remained similar at 1- and 6-months post-KP.2 vaccine dose, indicating sustained antigenic differences. Overall, neutralizing antibody titers against KP.2 increased in the majority of individuals after vaccination, although the magnitude of the increase varied among participants.

**Fig. 7.**
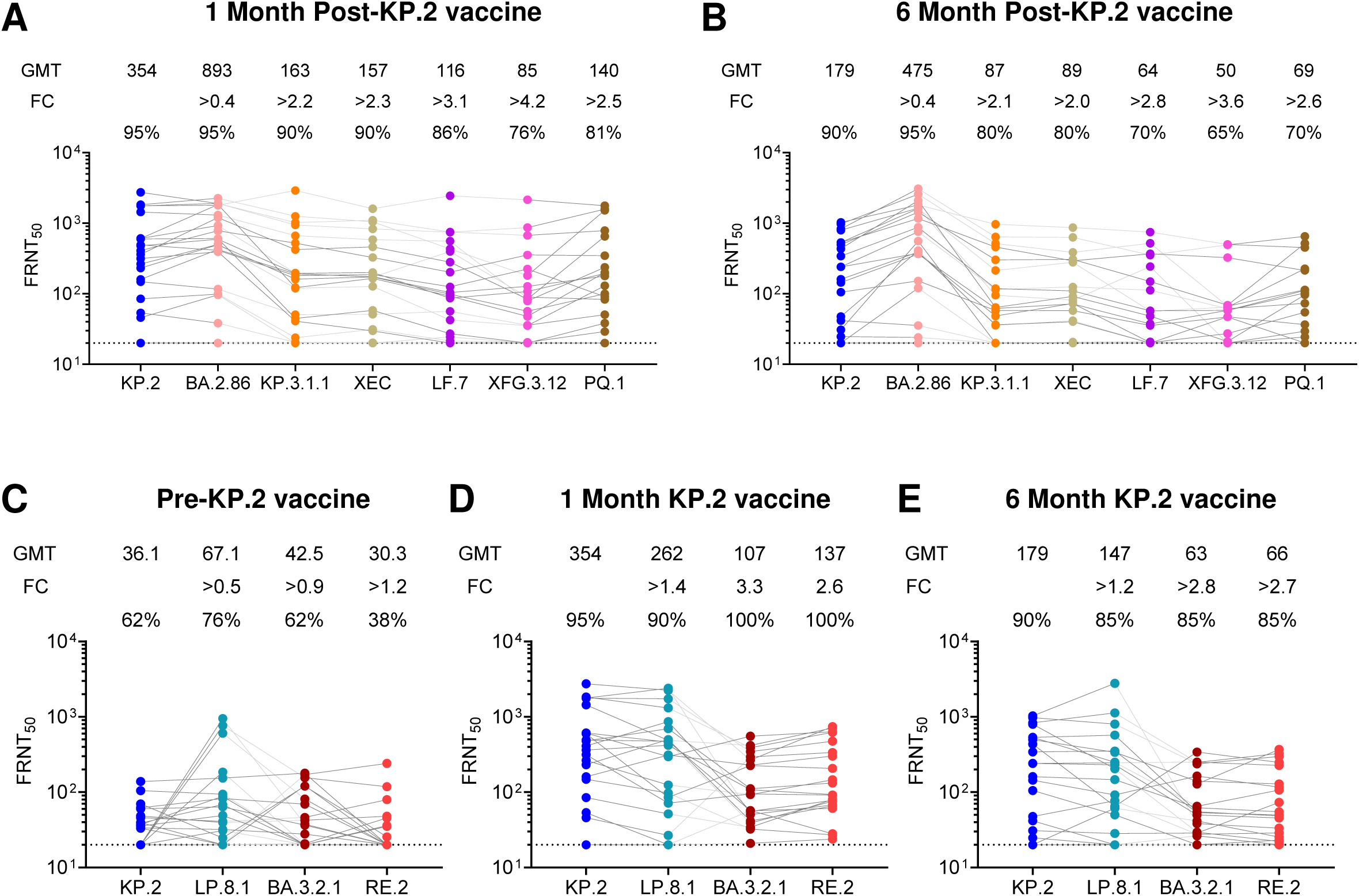
Neutralizing antibody responses against contemporary Omicron subvariants following KP.2 mRNA vaccination. **(A)** Neutralizing antibody titers (FRNT_50_) against KP.2 and emerging Omicron subvariants, including BA.2.86, KP.3.1.1, XEC, LF.7, XFG.3.12, and PQ.1 obtained for all 21 study participants are shown at **(A)** 1 month post-vaccination and **(B)** at 6 months post-vaccination. FRNT_50_ GMTs are indicated above each group. Fold change (FC) values are shown relative to KP.2. Each point represents an individual, and connecting lines show matched samples. (**C**) FRNT_50_ against KP.2, LP.8.1, BA.3.2.1 and RE.2 are shown for all 21 participants at (**C**) pre-, (D) 1 month post-, and (E) 6 months post-KP.2 vaccine dose.

The recent emergence of BA.3 subvariants further jeopardizes the antibody responses following the KP.2 vaccine dose (32). BA.3.2.1 was first detected in South Africa in November 2024 and is now found in circulation globally but still at relatively low levels (33). BA.3.2.1 and RE.2 consist of more than 50 mutaitons within the spike protein relative to KP.2 (34). We next evaluated the impact of these variants, along with the LP.8.1 which is the spike antigen encoded within the 2025 updated COVID vaccine (**Fig 3C-E**). Prior to the KP.2 vaccine dose, FRNT_50_ GMT was 67.1 for LP.8.1, 42.5 for BA.3.2.1, and 30.3 for RE.2 as compared to 36.1 for KP.2. The number of individuals with detectable neutralizing antibodies ranged from 62% for KP.2, 76% for LP.8.1, 62% for BA.3.2.1 and 38% for RE.2. Following the KP.2. vaccine dose, LP.8.1, BA.3.2.1 and RE.2 showed increased neutralizing antibody titers at 1 month post-vaccine dose and gradually declined through 6 months post-vaccine dose. Relative to KP.2, LP.8.1 showed a 3.9-fold increase in neutralizing antibodies between pre- and 1-month post-vaccine dose (FRNT_50_ 67.1 to 262) whereas BA.3.2.1 and RE.2 showed a 2.5- and 4.5-fold increase (FRNT_50_ 42.5 to 107 and FRNT_50_ 30.3 to 137), respectively. These findings suggests that the KP.2 vaccine does preferentially increases neutralizing antibody titers to the cognate KP.2 spike antigen as compared to either a closely related variant (LP.8.1) or more divergent variants (BA.3.2.1 and RE.2).

### LP.8.1 spike mRNA vaccination improves antibody neutralizing activity against LP.8.1 but not BA.3.2 variants

We next evaluated the neutralizing antibody titers in a small cohort of individuals (n = 6) that received the updated LP.8.1 spike mRNA vaccine dose (**Fig. 8**). Pre-LP.8.1 vaccine dose the FRNT_50_ GMT was 499 against WA1, 171 for BA.5, 119 for XBB.1.5, 108 for KP.2, 58 for LP.8.1, 49 for BA.3.2.1, and 45 for RE.2 (**Fig. 8A**). One month post-KP.2 vaccine dose, the FRNT_50_ GMT increased to 1015 for WA1, 524 for BA.5, 386 for XBB.1.5, 964 for KP.2, 692 for LP.8.1, 115 for BA.3.2.1, and 135 for RE.2 (**Fig. 8B**). The updated LP.8.1 vaccine dose showed significant increase in neutralizing antibody titers of 11.8-fold between pre- and 1 month post-vaccine dose as compared to WA1 (2.0-fold) and KP.2 (8.9-fold) (**Fig. 8C**). However, we observed a modest increase in neutralizing antibodies to BA.3.2.1 (2.3-fold) and RE.2 (3.0-fold). This suggests that the LP.8.1 vaccine dose preferentially boosts LP.8.1, and to a lesser extent KP.2, as compared to WA1, BA.3.21 and RE.2. Overall, these results indicate a clear enhancement of cross-reactive neutralizing activity following both the KP.2 and LP.8.1 vaccine dose. These findings demonstrate that boosting with the KP.2 or LP.8.1 spike mRNA vaccine enhances both homologous and cross-reactive neutralizing antibody responses, increasing neutralization potency against LP.8.1, KP.2, and to a lesser extent divergent Omicron subvariants.

**Fig. 8.**
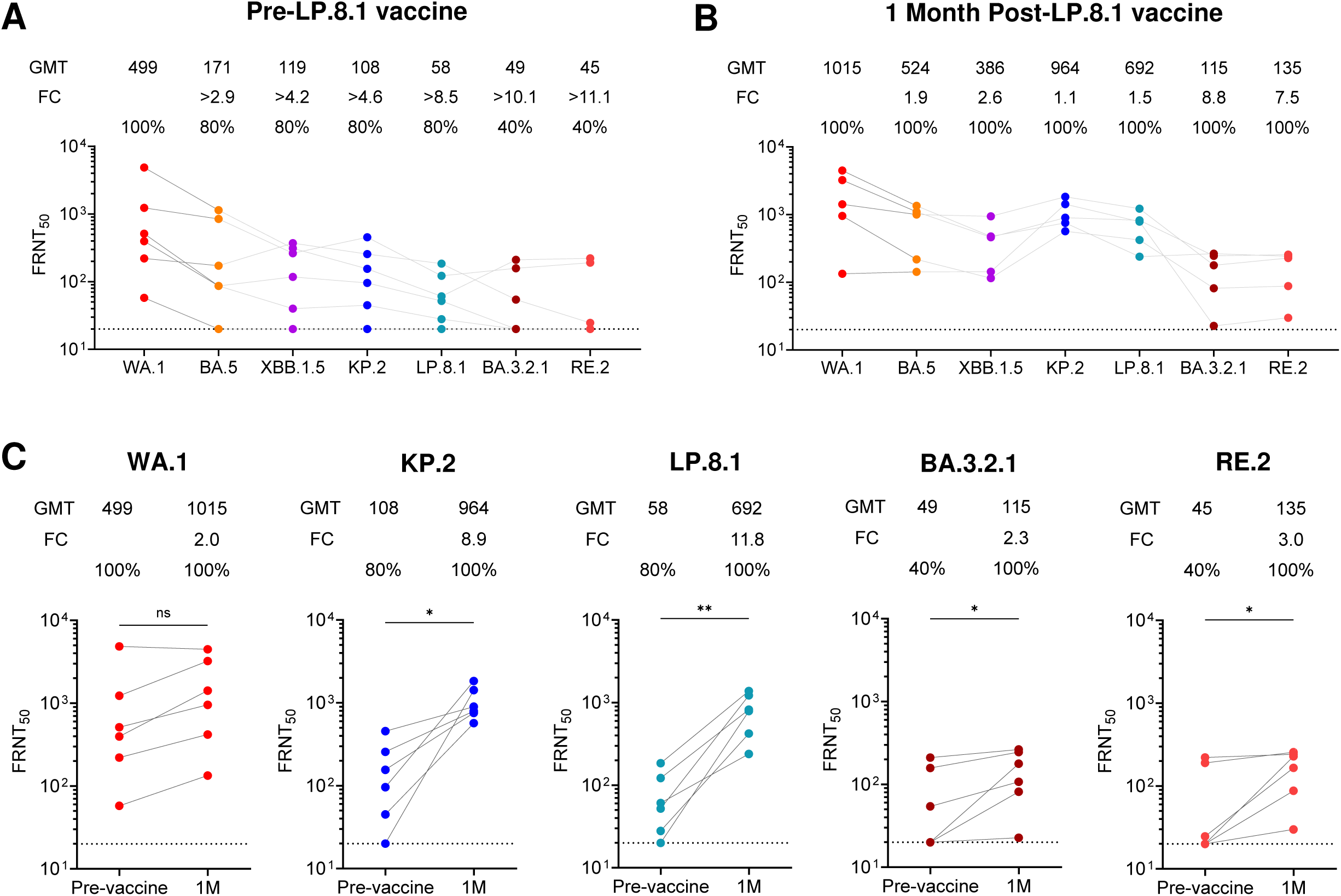
Neutralizing antibody responses following LP.8.1 mRNA vaccination in a subset of individuals. Neutralizing antibody titers (FRNT_50_) were measured against WA1, BA.5, XBB.1.5, KP.2, LP.8.1, BA.3.2.1, and RE.2 variants for LP.8.1 vaccination study (n = 5). **(A)** Shows FRNT_50_ titers obtained prior to LP.8.1 vaccine dose and **(B)** shows FRNT_50_ titers at 1 month following LP.8.1 vaccination. FRNT_50_ GMTs are indicated above each group along with the FC of the neutralization GMT against WA1 and the Omicron subvariants. **(C)** Shows a paired comparison of FRNT_50_ titers for WA1, KP.2, LP.8.1, BA.3.2.1 and RE.2 between pre- and 1 month post-vaccination time points. Each point represents an individual. FRNT_50_ GMTs are indicated above each group along with the FC of the neutralization GMT for pre-vaccination and 1 month following LP.8.1 vacine dose. Connected lines show paired samples of each individual. Statistical comparisons were performed using non-parametric t-test or Wilcoxon matched-pairs signed rank test. Here, ns = not significant, *p < 0.05, **p<0.01.

## DISCUSSION

In this study, we show that immunization with a KP.2 spike based SARS-CoV-2 mRNA vaccine induces robust and durable antibody responses with broad cross-reactivity against both ancestral and recently circulating Omicron variants. The KP.2 vaccine increased neutralizing activity against the matched KP.2 strain as well as antigenically distinct variants, including WA1, and BA.5, XBB.1.5. Although neutralizing titers declined over time, antibody responses remained detectable through six months, and neutralizing antibodies showed greater stability than binding antibodies. These findings indicate that updated vaccines based on contemporary Omicron lineages can enhance both the magnitude and persistence of humoral immunity. The antigenic distance between KP.2 and earlier SARS-CoV-2 variants is reflected in both our phylogenetic and structural analyses, which show accumulation of mutations in key regions of the spike protein. Despite this divergence, vaccination with KP.2 elicited cross-reactive antibody responses capable of recognizing multiple variants. The observed increase in neutralizing activity against WA1 following KP.2 vaccination suggests that conserved epitopes continue to contribute to the antibody response. This observation is consistent with prior studies reported by us and others that additional variant adapted vaccine doses can maintain or enhance responses to earlier strains while expanding coverage to new variants (11, 13, 24, 35-38).

A key observation from this study is the durability of antibody responses following KP.2 vaccination. Using a power law decay model, we estimated sustained half-lives for spike-specific IgG responses, particularly for WA1. The WA1 binding titers showed early decline relative to neutralizing antibody titers, suggesting the establishment of long-lived plasma cell responses. The strong correlation between binding and neutralizing antibody levels supports the use of binding assays as a surrogate for functional activity in longitudinal studies. The analysis of IgG subclasses revealed a response dominated by IgG1, with modest contributions from IgG2 and IgG4 subclasses. The increase in IgG4 following KP.2 vaccination is notable, particularly in the context of repeated exposure to SARS-CoV-2 antigens through vaccination is consistent with observations for the XBB.1.5 vaccine dose (11, 37, 39, 40). IgG4 antibodies have distinct structural and functional properties, including reduced ability to engage Fc-mediated effector functions (41, 42). While the biological significance of this shift remains unclear, it may reflect ongoing maturation of the antibody response under conditions of repeated antigen exposure (43). Further studies are needed to determine whether this subclass distribution influences protective immunity or affects responses to subsequent vaccine doses.

Using the spike depletion assay, the majority of spike-binding antibodies after KP.2 vaccination were cross-reactive between WA1 and KP.2, indicating that prior immune history still plays a dominant role in shaping the response. This is similar to our recent observations following the XBB.1.5 vaccine dose (11). This limited proportion of KP.2 specific antibodies suggests that vaccination primarily recalls memory B cells targeting conserved epitopes rather than generating a large pool of KP.2 variant specific immune responses. This pattern is consistent with immune imprinting, where earlier exposures influence the hierarchy of antibody specificities following subsequent immunization (31). While cross-reactive antibodies may provide broader protection, they may also limit the ability to fully adapt to antigenically distant variants. Therefore, future studies need to be carried out to define the vaccination strategies and vaccine antigen selection to overcome immune imprinting.

We further evaluated the breadth of neutralizing activity against a panel of recently circulating Omicron subvariants. KP.2 vaccination elicited measurable neutralization against all variants tested, including BA.2.86 spike that comprises 34 mutations and distantly related BA.3.2 variants consisting 52 spike mutations alon with other more recently emerged lineages (44, 45). However, reduced neutralization was observed for certain variants, particularly XFG.3.12 followed by LF.7, indicating ongoing antigenic drift. These reductions, while modest, highlight the continued evolution of the spike protein and its impact on antibody recognition. The persistence of detectable neutralizing activity at six months suggests that protection against severe disease may be maintained, even as susceptibility to infection may increase over time. The analysis of a few individuals who received the recently updated LP.8.1 spike mRNA vaccine dose provides additional insight into how updated immunogens shape the antibody response. Vaccination with LP.8.1 improved neutralizing activity against the matched variant while maintaining cross-reactivity with earlier strains. Interestingly, the neutralizing antibody titers were comparable between WA1 and KP.2 variant in these individuals, but limited increased titers were obsevd against more diverse Omicron variants BA.3.2.1 and RE.2, suggesting the involvement of KP.2 variant specific recall responses. This observation supports the concept that sequential updates to vaccine antigens can incrementally broaden the antibody repertoire, although the extent of this effect may depend on the degree of antigenic difference and prior exposure history.

This study has several limitations that are woth noting. The cohort size is modest, which may limit the ability to detect smaller effects and increases variability in subgroup analyses. The follow-up period is restricted to six months, and longer-term studies are needed to define the durability of responses beyond this time frame. We did not assess T cell responses, which are likely to contribute to protection against severe disease. In addition, we did not directly analyze the epitope specificity or clonal composition of memory B cells, which would provide a more detailed understanding of the mechanisms underlying cross-reactivity and imprinting. The impact of prior infection history was not fully resolved and may contribute to heterogeneity in the observed responses.

In summary, KP.2 spike mRNA vaccine dose induces durable and broadly cross-reactive antibody responses against SARS-CoV-2, including recently circulating Omicron variants. At the same time, reduced neutralization of certain variants and the dominance of cross-reactive antibodies highlight the challenges posed by ongoing viral evolution and immune imprinting. These findings support continued monitoring of antigenic changes and periodic updating of vaccine formulations to maintain effective protection against emerging variants.

## MATERIALS AND METHODS

### Cells and Viruses

VeroE6-TMPRSS2 cells were generated and cultured as previously described (46). nCoV/USA_WA1/2020 (WA1), closely resembling the original Wuhan strain, was propagated from an infectious SARS-CoV-2 clone as previously described (47). Omicron subvariants were isolated from residual nasal swabs. BA.5 isolate (EPI_ISL_13512579), XBB.1.5 (EPI_ISL_16026423), and KP.2 (EPI_ISL_18993067) was previously described (11, 13). BA.2.86, KP.3.1.1, XEC, LF.7, XFG.3.12, PQ.1, LP.8.1, BA.3.2.1 and RE.2 were provided by the Centers for Disease Control and Prevention (CDC) (**Table S2**). All variants were propagated once in VeroE6-TMPRSS2 cells to generate working stocks. Viruses were deep sequenced and confirmed as previously described (48).

### Focus Reduction Neutralization Test (FRNT)

Live virus based focus reduction neutralization test (FRNT) assays were carried out following previously established protocols with minor adjustments (46, 48, 49). Serum samples were tested in duplicate and subjected to eight serial three-fold dilutions in DMEM, starting at an initial dilution of 1:10. Each diluted serum sample was combined with an equal volume of live SARS-CoV-2 variant virus, corresponding to approximately 100 to 200 infectious foci per well. The serum virus mixtures were incubated for 1 hour at 37°C in round-bottom 96-well plates to allow antibody virus interactions. Following incubation, the mixtures were transferred onto monolayers of VeroE6-TMPRSS2 cells and incubated for an additional hour at 37°C. After this adsorption step, the inoculum was removed and replaced with 100 µl of prewarmed 0.85% methylcellulose overlay medium in each well to restrict viral spread. Plates were then incubated at 37°C for 18 to 40 hours depending on the replication kinetics of the specific viral variant. At the end of the incubation period, the overlay medium was carefully removed, and cells were washed with PBS and fixed using 2% paraformaldehyde for 30 minutes. Fixed cells were washed twice with PBS and subsequently permeabilized for at least 20 minutes using permeabilization buffer. Viral foci were detected by incubating the cells overnight at 4°C with an Alexa Fluor 647 conjugated anti-SARS-CoV-2 spike monoclonal antibody (CR3022-AF647). Plates were washed twice with PBS before imaging and quantification using an ELISPOT reader (CTL Analyzer).

### Spike protein-binding assay

Spike protein specific IgG antibodies against SARS-CoV-2 and Omicron subvariants were measured using an electrochemiluminescence multiplex immunoassay performed on the Meso Scale Discovery platform as describe previously (11). Measurements were carried out with the V-PLEX SARS-CoV-2 Key Variant Spike Panel 39 kit (catalog no. K15738U for IgG) together with internally prepared reagents for subclass detection. Serum samples were diluted 1:5000 before being added to the assay plates. Plates supplied with the kit were first incubated with blocking buffer A, provided by the manufacturer, for 30 minutes. For determination of IgG subclasses, detection antibodies specific for each subclass were applied at a final dilution of 1:3000 (50 µl per well). These included mAbTech anti-IgG1-sulfo, Southern Biotech anti-IgG2-sulfo, mAbTech anti-IgG3-sulfo, and mAbTech anti-IgG4-sulfo. Signals were acquired and processed using Discovery Workbench software and further analyzed in GraphPad Prism version 10.1.2. Antibody levels were quantified by comparison with standard control curves and are reported as arbitrary units per milliliter for total IgG or as micrograms per milliliter for IgG subclass measurements.

### Spike protein trimer S2P antigens

For serum depletion studies, stabilized SARS-CoV-2 spike trimer proteins (spike-6P) representing the WA.1 strain and the KP.2 variant were produced in mammalian cells. Expression constructs encoding WA.1 spike-6P and KP.2 spike-6P were introduced into Expi293F cells as described previously (50). Transfections were carried out using the ExpiFectamine 293 system (Thermo Fisher, catalog no. A14524) following the manufacturer’s recommended protocol. Expi293F cultures were maintained at a density of approximately 2.5 x 10⁶ cells per ml at the time of transfection. Following incubation for 4 to 5 days to allow protein expression, cell cultures were clarified by centrifugation at 4,000 x g for 20 minutes at room temperature. The resulting supernatant containing secreted spike protein was collected and passed through a filter before purification. The clarified supernatant was then applied to a pre-equilibrated Excel Ni-NTA affinity column. After sample loading, the column was washed with PBS containing 25 mM imidazole to remove non-specific proteins, and the bound spike protein was subsequently eluted using PBS supplemented with 250 mM imidazole. Eluted fractions were concentrated and further purified by size-exclusion chromatography on a Superose 6 Increase 10/300 column using PBS as the running buffer. The spike protein eluted as a trimeric species, and the corresponding peak fractions were collected. Protein integrity and purity were assessed by SDS-PAGE and by negative-stain electron microscopy.

### WA1 and KP.2 spike protein depletion assay

Depletion of WA1 or KP.2 spike specific antibodies from plasma was performed following an established protocol described early with minor adjustments (11). In brief, recombinant SARS-CoV-2 spike proteins corresponding to WA1 spike-6p or KP.2 spike-6p or the WA1 RBD and KP.2 RBD were chemically conjugated to paramagnetic M-270 epoxy Dynabeads using the Dynabeads Antibody Coupling Kit (Thermo Fisher Scientific, catalog no. 14311D), following the manufacturer’s guidelines for coupling 60 mg of beads. In separate tubes, 30 mg of beads were prepared using 20 µg of Spike protein per mg of beads. After coupling, beads were suspended at a concentration of 10 mg/ml in storage buffer at 4°C. Immediately before use, Spike-coupled beads were washed once for 5 min in PBS with 0.1% BSA and then resuspended in DPBS. Plasma samples were added to beads at a ratio of 1:50 (v/v) and gently mixed for 1 h at room temperature using a rotating mixer. Depleted plasma were separated from beads with a magnet tube rack and transferred to a fresh tube that contained Spike-coupled beads equal in amount to the first depletion, which had been separated from the storage solution. Samples were incubated again for 1 h at room temperature and then magnetically separated from beads yielding Spike-depleted plasma diluted 1:50 in DPBS. Samples were then ready to use in MSD or RBD binding assays, as described above. Percentage reduction was then calculated from the fold change between the pre-depletion and post-depletion samples.

### WA1 and KP.2 RBD ELISA assay

Recombinant SARS-CoV-2 RBDs WA1, and KP.2 were coated on separate Nunc MaxiSorp plates at a concentration of 1 μg/mL in 100 uL phosphate-buffered saline (PBS) at 4°C overnight. Plates were blocked for 1.5 hours at room temperature in PBS/0.05%Tween/1% BSA (ELISA buffer). Samples were serially diluted 1:2 in dilution buffer (PBS-1% BSA-0.05% Tween-20) starting at a dilution of 1:500. 100 μL of each dilution was added and incubated for 90 minutes at room temperature. 100 uL of horseradish peroxidase-conjugated anti-human-IgG-HRP (Jackson Immuno Research) secondary antibody, diluted 1 to 3,000 in ELISA buffer, was added and incubated for 60 minutes at room temperature. Development was performed using 0.4 mg/mL o-phenylenediamine substrate (Sigma) in 0.05 M phosphate-citrate buffer pH 5.0, supplemented with 0.012% hydrogen peroxide before use. Reactions were stopped with 1 M HCl and absorbance was measured at 490 nm. Between each step, samples were washed three times with 300 uL of PBS-0.05% Tween. Prior to development, plates were additionally washed once with 300 uL of PBS.

### Phylogenetic analysis

SARS-CoV-2 spike protein tree based on amino acid sequence and generated in Geneious Pro Software (Biomatters Ltd) using Blosum 62 scoring matrix, open gap penalty of 12 and gap extension penalty of 3, the Jukes-Cantor genetic distance model, and Neighbor-Joining tree building method. SARS-CoV-2 spike protein IDs: Wuhan-Hu-1 (P0DTC2); Alpha (B.1.1.7, XIF92171.1); Beta (B.1.351, XIF12677.1); Delta (B.1.617.2, XIF92419.1); Omicron BA.1.1 (XHH56286.1); Omicron BA.2 (XCA50803.1); Omicron BA.5 (UOZ45804.1); Omicron XBB1.5 (UZG29433.1); Omicron JN.1 (XHE63187.1); Omicron KP.2 (XIF87639.1); Omicron KP.3 (XIF89987.1); Omicron XEC (XII33585.1).

### Statistical Analysis

Mixed effects exponential, power law, and biphasic decay models implemented in Monolix (Monolix, Lixoft) were used to analyze the waning of the antibody response. The exponential and power law models were formulated as ordinary differential equations, 𝑑𝐴𝑏/𝑑𝑡=−𝑘·𝐴𝑏 and 𝑑𝐴𝑏/𝑑𝑡=−𝑘/𝑡·𝐴𝑏, respectively, where parameters 𝑘 are corresponding decay rates, with antibody titers at day 18 to be log-normally distributed and with log-normal multiplicative error. The biphasic decay model assumed two phases, each characterized by exponential decay. The estimation of the population parameters was performed using the Stochastic Approximation Expectation-Maximization (SAEM) algorithm. Half-lives were calculated as ln(2)/𝑘 for the exponential model and for both phases of the biphasic model, and as *T*(2^𝑘^-1), where *T* is the time at which the half-life is measured for the power-law model (*T* = 120 days). Antibody neutralization was quantified by counting the number of foci for each sample using the Viridot program(51). The neutralization titers were calculated as follows: 1 – (ratio of the mean number of foci in the presence of sera and foci at the highest dilution of the respective sera sample). Each sample was tested in duplicate. The FRNT_50_ titers were interpolated using a 4-parameter nonlinear regression in GraphPad Prism 10.1.2. Samples that do not neutralize at the limit of detection at 50% are plotted at 20 and used for geometric mean and fold-change calculations. Kruskal-Wallis test with Dunn’s multiple comparison test was used to analyze difference in neutralization and IgG antibody binding titers. A Wilcoxon signed-rank test was performed to compare IgG subclass titers between paired samples from pre- and post-1 month vaccination. Pearson’s rank test was used for the correlation analysis. Correlation analyses were done by log transforming spike protein binding titers or neutralization titers, followed by linear regression analysis. The *p* values are reported in each figure.

## ACKNOWLEDGEMENTS

We thank our collaborators at the Centers for Disease Control and Prevention (CDC) Jennifer L. Harcourt, Clinton R. Paden, Meredith Gardner, Heather Hicks, Azaibi Tamin, and Peter Cook for providing SARS-CoV-2 variants. We also thank Natalie J Thornburg and the CDC Traveler’s Genomic Surveillance Program (TGS), who provide us with the patient swabs used to generate the clinical isolates. Funders played no role in the design and conduct of the study; collection, management, analysis, and interpretation of the data; preparation, review, or approval of the manuscript; and decision to submit the manuscript for publication.

## AUTHOR CONTRIBUTIONS

S.K., J.W, A.M., V.Z., A.P, E.A.O and M.S.S. contributed to the analysis, interpretation of data and writing the manuscript. L.L and J.V.V isolated, propagated, and sequenced the viruses. A.P purified recombinant SARS-CoV-2 protein. V.D.M performed phylogenetic analysis on SARS-CoV-2 strains. L.L. and M.L.E. performed in vitro neutralization experiments. V.Z. performed the statistical analysis. J.V.V., L.L., and M.L.E. processed the samples for PBMCs isolation. D.J. and M.L.E. performed binding assays. S.K., D.J., and M.L.E. performed IgG subclass binding assays and calculations. J.Z.A.F, S.T.W, R.R.P., J.I, K.B, S.E., and N.R. provided clinical support for the study and contributed to sample collection. J.W. and M.S.S. contributed to the conception and design of the work and the writing and approval of the final manuscript.

## DATA AVAILABILITY

All data associated with the paper are available in the main text or the supplementary materials. Requests for SARS-CoV-2 isoaltes should be addressed to Mehul S. Suthar and can be made available to academic researchers without an MTA but with appropriate laboratory containment.

## ETHICS APPROVAL

Peripheral blood samples from donors were collected at Emory University (Emory National Primate Research Center, Emory Hope Clinic, and Emory Children’s Center). Adults ≥18 years old who met eligibility criteria under these protocols and provided informed consent were enrolled. All patient samples were de-identified prior to inclusion in the study. Sample collection and processing were performed with approval from the University Institutional Review Board (#00002061, #00058271 and #00022371). For all of the donors, samples were collected before the KP.2 vaccine dose, 1 month, 3 months, and 6 months after the KP.2 vaccine dose. For LP.8.1, samples were collected prior to and 1 month after the vaccine dose. Blood samples were processed to obtain plasma and peripheral mononuclear cells.

## Supplemnetal Material

Figures S1

Table S1 and S2

## SUPPLEMENTARY FIGURE LEGENDS

**Fig. S1.**
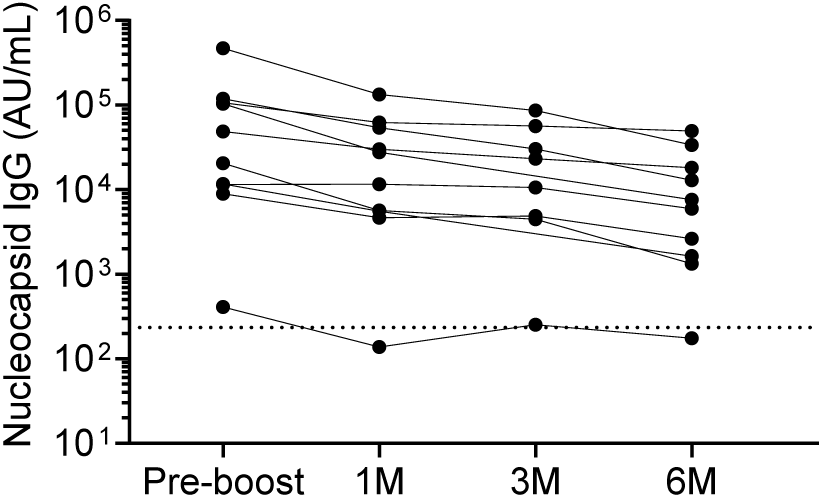
Nucleocapsid-specific antibody responses during the six-month follow-up after KP.2 mRNA vaccination. IgG antibody binding to the SARS-CoV-2 nucleocapsid (N) protein measured by electrochemiluminescent immunoassay across the longitudinal sampling period. Antibody levels are shown for a subset of participants (n = 10) included for durability analysis at baseline prior to KP.2 vaccination (pre-vaccination) and at 1 month (1M), 3 months (3M), and 6 months (6M) after vaccination. Each symbol represents an individual participant and lines connect longitudinal samples from the same individual. Antibody concentrations are reported as arbitrary units per milliliter (AU/mL) based on normalization to a standard reference curve.

